# A rhinovirus vaccine evokes T cell mediated cross-reactive immunity in a preclinical model of rhinovirus infection

**DOI:** 10.64898/2026.01.06.697896

**Authors:** Stephen M Shaw, Chloe J Pyle, Neeta D Patel, Michael R Edwards, Ziyin Wang, John S Tregoning, Patricia Londoño-Hayes, Paul A Hamblin, Richard P Butt, Sebastian L Johnston

**Author notes:** **Corresponding authors:** Stephen M Shaw and Sebastian L Johnston. These authors contributed equally (Stephen M Shaw, Chloe J Pyle). **Author contribution statement** S.M.S and C.J.P. contributed equally. Study conception: S.M.S., P.A.H., R.P.B., S.L.J. Experimental design: S.M.S., C.J.P., N.D.P., M.R.E., J.S.T., P.L-H., R.P.B., S.L.J. Data acquisition: C.J.P., N.D.P., M.R.E., Z.W. Data interpretation and analysis: S.M.S., C.J.P., N.D.P., M.R.E., Z.W., J.S.T., P. L-H., S.L.J. Manuscript first draft: S.M.S. Manuscript revision and editing: S.M.S., C.J.P., P. L-H., S.L.J. All authors have read and approved the manuscript. **Funding declaration** This study was funded by Apollo Therapeutics Limited. **Competing interest declaration** S.M.S. and R.P.B. are employees of, and shareholders in, Apollo Therapeutics Limited. P.A.H. was an employee of Apollo Therapeutics Limited while engaged in the research project and is currently a consultant for Apollo Therapeutics Limited. P.L-H. is a consultant for Apollo Therapeutics Limited. C.J.P., N.D.P. and M.R.E. are employees of Virtus Respiratory Research Limited. M.R.E. and S.L.J. are shareholders in Virtus Respiratory Research Limited. J.S.T. and Z.W. have no competing interests to declare.

## Abstract

Rhinovirus (RV) infection is the most common cause of acute exacerbations of respiratory diseases such as asthma and chronic obstructive pulmonary disease. Vaccines are available for two less common respiratory viruses (influenza and respiratory syncytial viruses), but development of a RV vaccine to protect patients with chronic respiratory disease has been hampered by the large number of serologically distinct RV strains (∼180, grouped genetically into ∼80 A, ∼30 B and ∼70 C strains). We have explored the potential of the RV-A16 VP0 subunit vaccine in both CpG-adjuvanted protein subunit and mRNA formats. We demonstrate that both vaccine formats induce strong effector CD4 and CD8 T cell responses in the airways and lungs of immunized mice that rapidly expand following infection with a heterotypic RV strain and are associated with enhanced virus clearance. Adjuvanted VP0 protein subunit vaccination evoked strong germinal center reactions in the lymph nodes and markedly accelerated the production of neutralizing antibodies following heterotypic RV infection. In contrast, VP0 mRNA vaccination evoked limited germinal center responses and neutralizing antibody production. Th1 polarized immune responses were induced against a large panel of 20 heterologous RV strains representative of all RV-A, -B and -C strains. This study confirms the potential of RV VP0 vaccination as an approach to overcome the serological diversity of RV and describes the immune response evoked, including cross-strain cellular immunity and protection against heterotypic RV infection.

## Introduction

Rhinoviruses (RV) are the most common etiological agent of the common cold in humans. In healthy individuals the symptoms of RV infection are mild and mostly confined to the upper respiratory tract. However, in patients with chronic lung conditions, such as chronic obstructive pulmonary disease (COPD) or asthma, RV infection can lead to severe lower respiratory tract symptoms and cause acute exacerbations^1–3^ . RV infection has been associated with ∼80% of asthma exacerbations in children and ∼50-60% of COPD exacerbations in adults^4–10^ .

RVs are single-stranded positive-sense RNA viruses of the family *Picornaviridae*. RVs are divided genotypically into 3 species: RV-A, B and C, each of which includes many subtypes; serotypes from the RV-A and RV-C species are most often associated with exacerbations of asthma and COPD^11,12^ . There have been several attempts to develop a vaccine, primarily to protect patients with chronic lung diseases from RV-induced acute exacerbations^13–19^ . However, the large number of genotypically distinct subtypes (∼180) and high degree of inter-and intra-species antigenic diversity has hampered attempts to date.

We have previously reported the identification of highly conserved regions of cross-species homology located within the VP0 polyprotein of RV-A and RV-B virus types^20^ . VP0 is an immature RV capsid protein that gets cleaved into VP4 and VP2 during virion maturation, prior to egress from the host cell. An adjuvanted subunit vaccine candidate, based upon the VP0 protein of RV-A16 has been evaluated preclinically^20^. Mice immunized with recombinant RV-A16 VP0 formulated with CpG and Incomplete Freund’s Adjuvant (IFA), developed a Th1 polarized antiviral immune response, which reacted with VP0 from RV-A16 and cross-reacted with 3 other (2 RV-A and one RV-B) subtypes (RV-A1B, RV-B14 and RV-A29). Vaccination with RV-A16 VP0 also boosted RV-A1B-specific neutralizing antibody production and enhanced virus clearance from mice following subsequent intranasal infection with the heterologous RV-A1B isolate.

In this study, we further explored the potential of RV-A16 VP0 as a vaccine candidate and, given the current interest in mRNA platforms, compared the immune responses elicited by VP0 presented as an adjuvanted protein subunit vaccine or a mRNA vaccine. We used pre-clinical models to demonstrate that both vaccine platforms evoke Th1-biased, broadly cross-reactive anti-RV cellular and humoral immunity, associated with the expansion and maturation of B and T cells, formation of long-lasting tissue-resident memory T cells, and protection from subsequent infection with a heterologous subtype of RV.

## Results

### RV-A16 rVP0 immunization generates strong antigen-specific humoral and cellular responses accompanied by mobilisation of T and B cell populations in lymphoid tissue

Mice were immunized with RV-A16 rVP0 adjuvanted with CpG and IFA (RV-A16 rVP0 vaccine) or with adjuvant only (mock immunization control), on two occasions, 21 days apart (Fig. 1A). A single immunization with RV-A16 rVP0 vaccine was sufficient to generate a strong VP0-specific IgG serum response in all immunized mice and a second vaccination further boosted these responses (Fig. 1B). Mock immunization with adjuvant only did not induce an antigen specific response.

**Figure 1.**
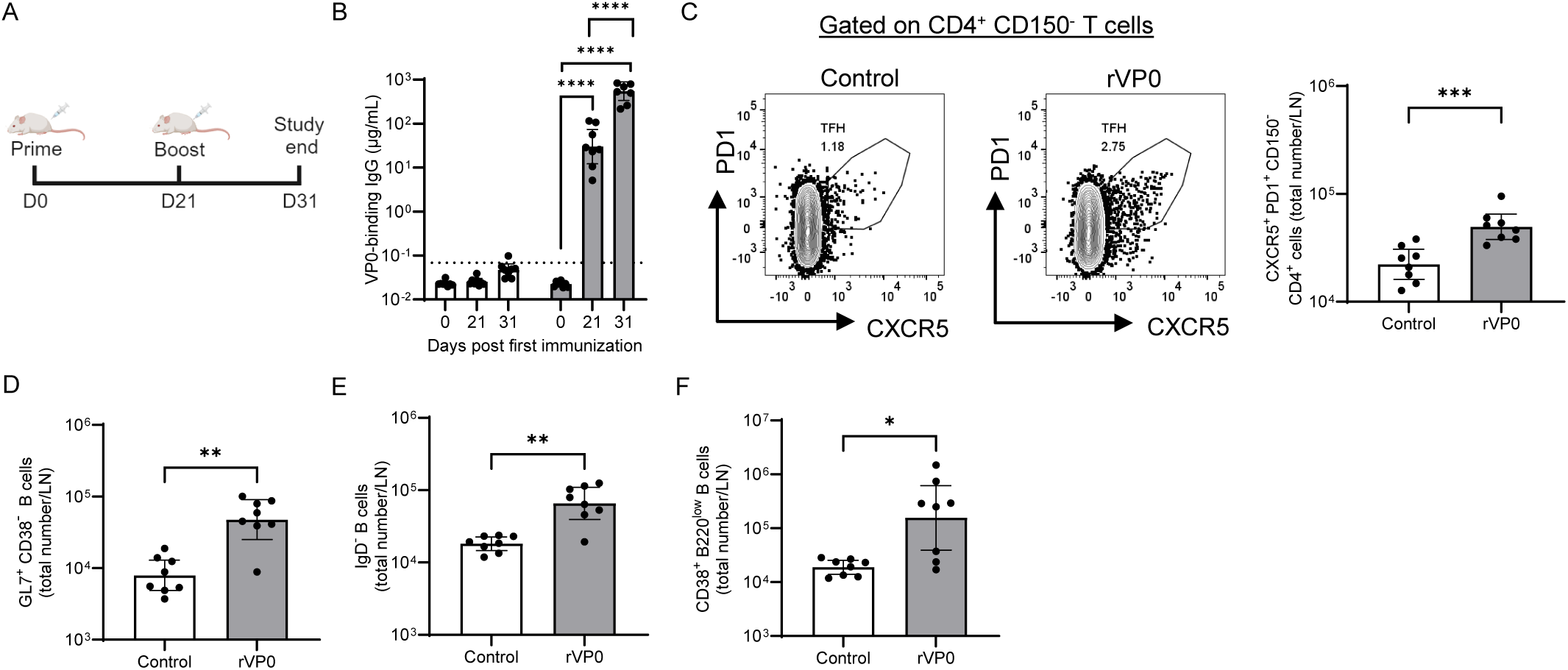
RV-A16 VP0 protein immunization stimulates production of RV-A16 VP0-specific IgG and inguinal lymph node T and B cell responses. (A). Mice were immunized twice with adjuvant only control (open bars) or recombinant RV-A16 VP0 adjuvanted with CpG and IFA (grey bars) 21 days apart. 10 days after the final immunization lymph nodes were harvested. Blood was taken on study days Day 0, 21 and 31. Each point represents an individual mouse. (B). Serum concentrations of RV-A16 VP0-binding total IgG at each timepoint. Dotted line indicates lower limit of quantification (LLOQ), defined as half of the last point on the standard curve. ****p<0.0001 (ANOVA). The abundance of inguinal lymph node T_FH_ cells (C), germinal center B cells (D), class-switched B cells (E) and plasma cells (F) were determined by flow cytometry. *** p<0.001, ** p<0.01, * p<0.05 (Mann-Witney). Graphs show geomeans with 95% CI. Figure 1A created using BioRender.com.

Analysis of inguinal lymph nodes (LN) from vaccinated mice after the second immunisation showed that this strong humoral response was accompanied by differentiation of T and B cell populations in lymphoid tissue. Ten days after the second immunisation (study day 31), there were significantly more T follicular helper cells (Fig. 1C), germinal centre B cells (Fig. 1D), class switched B cells (Fig. 1E) and plasma cells (Fig. 1F) in mice immunized with RV-A16 rVP0 than in mice immunized with adjuvant only control.

### Heterologous RV infection evokes rapid anamnestic neutralizing antibody responses and cross-reactive T cell responses in mice previously vaccinated with RV-A16 VP0 protein

To assess whether RV infection could recall an immune response in mice previously immunized with the RV-A16 rVP0 vaccine, the antibody response was examined in vaccinated animals after intranasal challenge with a heterologous RV-A1B virus isolate (Fig. 2A). In parallel, groups of vaccinated animals were given a mock-challenge of PBS to serve as sham-infected controls. On days 21 and 49 following immunization, a strong antibody response was present in sera from mice immunized with RV-A16 rVP0 vaccine, but not in sera from mice immunized with adjuvant only (Fig. 2B). RV-A1B infection did not generate a significant change in binding antibody response against VP0 in mock-immunized mice, neither did it have a significant impact on the titer of the VP0-IgG response in RV-A16 rVP0 vaccinated mice (Fig. 2B).

**Figure 2.**
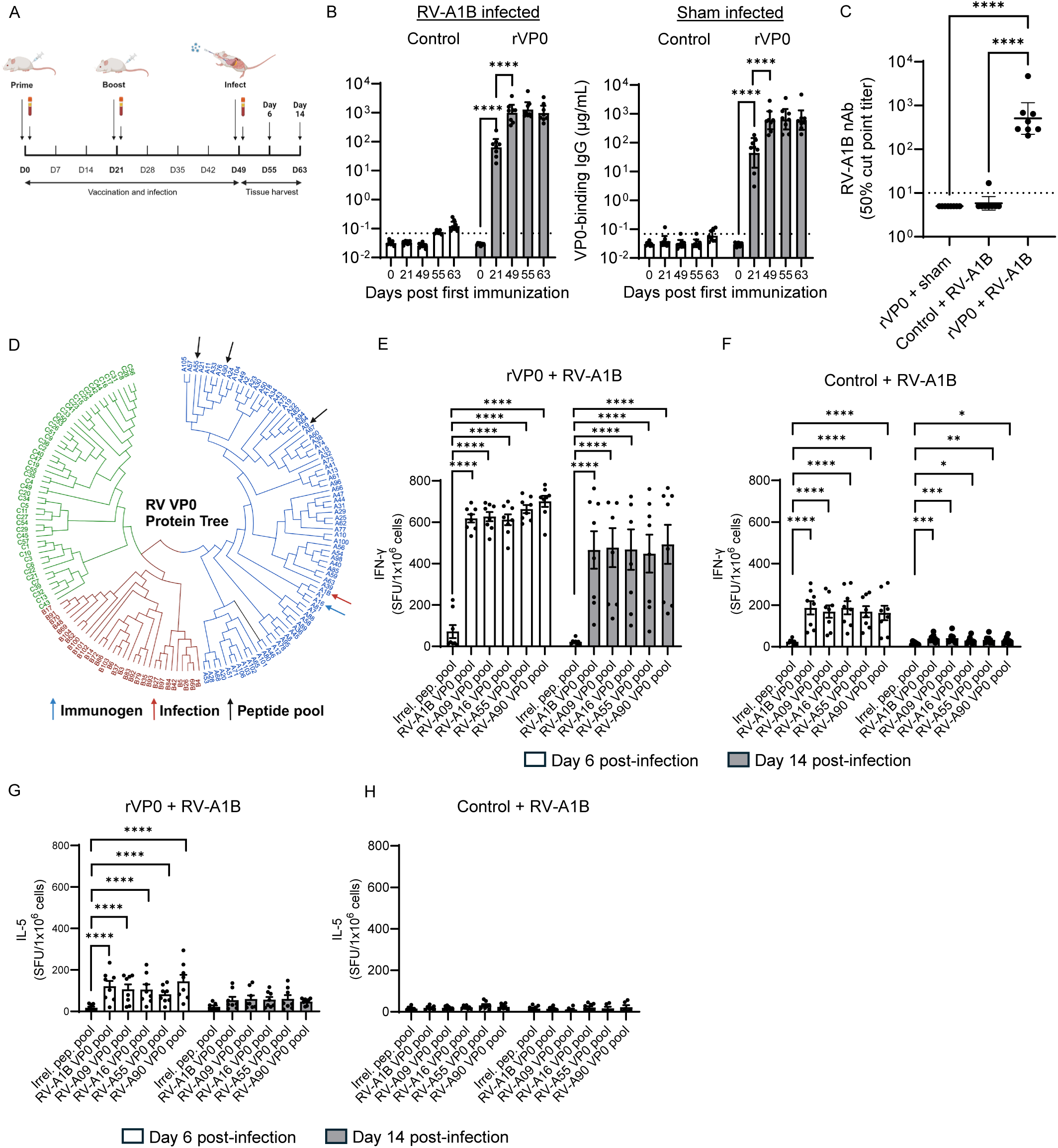
RV-A16 VP0 protein immunization accelerates production of neutralizing antibodies to heterotypic RV-A strain infection and creates cross-reactive cellular anti-RV immunity. (A). Mice were immunized twice with adjuvant only control or recombinant RV-A16 VP0 adjuvanted with CpG and IFA 21 days apart. 28 days after the final immunization mice were infected intranasally with live RV-A1B. Blood was taken on study days Day 0, 21, 49, 55 (6 days post infection) and 63 (14 days post-infection). The concentration of serum RV-A16 VP0-binding total IgG was determined at each timepoint in sham or RV-A1B infected animals (B) (dotted line indicates LLOQ). The titer of RV-A1B neutralizing serum antibodies was determined on Day 6 post-infection (C), dotted line indicates LLOQ (defined as half of the first dilution factor). Spleens were harvested on study days 55 (Day 6 post-infection) and Day 63 (Day 14 post-infection), splenocytes prepared and stimulated with peptide pools corresponding to full-length VP0 (15-mers with 11 amino acid overlap) from different RV-A strains (D) or a negative control irrelevant peptide pool (human HLA class I Ig-like C1 type domain). IFN-γ or IL-5 secreting cells from recombinant RV-A16 VP0 (E & F) immunized mice or controls (G & H) or were quantified by ELISpot. **** p<0.0001, *** p<0.001, ** p<0.01, * p<0.05 (ANOVA). Graphs on log scale show geomeans with 95% CI, those on a linear scale show mean with SEM. Each point represents an individual mouse. Figure 2A was created using BioRender.com.

In contrast, clear differences were observed among study groups with regards to the magnitude of the heterologous neutralizing antibody (nAb) response triggered by RV infection. When the titer of RV-A1B neutralizing antibodies was assessed on Day 6 after challenge a rapid and significant increase in nAb, suggestive of an anamnestic response, was observed in RV-A16 rVP0 immunized mice, as compared to adjuvant only immunized mice, or to mice previously immunized with RV-A16 rVP0 vaccine subjected to a sham-challenge (Fig. 2C). These findings suggest that in RV-A16 rVP0 vaccinated animals that respiratory infection, even with a heterologous RV subtype, can recall rapid antibody responses with cross-neutralizing activity.

We have previously shown that immunization with RV-A16 rVP0 induces cross-reactive T cell responses against three other (2 RV-A and one RV-B) subtypes^20^ . In the current study, we assessed the impact of RV-A16 rVP0 immunization and heterotypic RV-A1B infection on these cellular responses to other RV-A subtypes. Splenocytes isolated on Day 6 and Day 14 post infection were stimulated with peptide pools corresponding to full-length VP0 from diverse RV-A subtypes including RV-A16 (vaccine homologous) and RV-A1B (heterologous challenge), which are closely related, but also more distantly related (RV-A09) and very distantly related (RV-A90 and RV-A55) subtypes (Fig. 2D), or an irrelevant peptide pool as negative control. To determine the predominant T helper (Th) phenotype of responding splenocytes, both IFN-γ and IL-5 ELISpot analysis was performed.

Strikingly, on Day 6 after RV-A1B infection, all RV-A peptide pools tested induced strong and durable IFN-γ response from splenocytes isolated from RV-A16 rVP0 vaccinated mice, with strong responses persisting to Day 14 (Fig. 2E). RV-A1B infection also triggered a modest IFN-γ response in splenocytes from adjuvant only immunized mice against all the RV-A VP0 peptide pools tested (Fig. 2F). However, this response was more transient (i.e. almost down to irrelevant peptide control levels by day 14 post-infection) and of lower magnitude than in RV-A16 rVP0 vaccinated mice, hence likely reflective of primary T cell activation caused by infection.

Following RV-A1B infection, modest, transient IL-5 responses were observed in splenocyte cultures from RV-A16 rVP0 vaccinated mice against all of the VP0 peptide pools tested (Fig. 2G), but not the irrelevant peptide pool. These responses were greater from splenocytes isolated on day 6 compared to day 14. No IL-5 secretion was observed in splenocytes from adjuvant-only immunized controls (Fig. 2H). Both on day 6 and day 14 post infection, spot counts of IL-5 secretion by splenocytes from RV-A16 rVP0 immunized mice were 5-10-times lower than those for IFN-γ secretion (Fig. 2E vs Fig. 2G), indicating a strong Th1 and modest Th2 induction following RV-A16 rVP0-vaccination.

### RV-A16 VP0 mRNA vaccine evokes a weak antibody binding and serum neutralizing response that cannot be effectively recalled by subsequent RV infection

Progress in the development and use of mRNA vaccine technologies has accelerated over last decade. Having established the immunogenicity of RV-A16 VP0 protein, we next assessed immunization using a RV-A16 VP0 mRNA vaccine.

To demonstrate that RV-A16 VP0 mRNA can drive expression of full length VP0 protein in mammalian cells, we transfected HEK293 cells and determined the expression of VP0 over time (Fig. S1). Expression of VP0 protein peaked at 48 hours after transfection and declined thereafter, consistent with functional translation of the mRNA.

To study the immune response to RV16 VP0 when delivered as an mRNA vaccine, mice were immunized on days 0 and 21 with RV-A16 VP0 mRNA formulated with in vivo-jetRNA®, or with in vivo-jetRNA® alone, as mock-immunization control. Four weeks after the second vaccination, immunized animals were split into groups and challenged by intranasal infection with a heterologous RV-A1B isolate, or with PBS as a sham-infection control. As with the rVP0 protein vaccine, serum IgG responses to VP0 were detectable on day 21 after primary mRNA vaccination and titers increased (day 49) following a second immunization (Fig. 3A). However, the response was approximately 2 orders of magnitude lower than that generated by vaccination with RV-16 rVP0 protein vaccine (Fig. 2B vs 3A). Moreover, subsequent intranasal infection with RV-A1B did not have a discernible effect on the VP0-specific antibody titers raised by RV-A16 VP0 mRNA (day 63), indicating that overall, the mRNA vaccine delivered with the formulation and timing used in this study was not as effective at generating antibody responses as the protein vaccine platform. In these mice, consistent with low levels of antigen-specific IgG, no indication of cellular mobilization was found in inguinal lymph nodes on day 10 after a second immunization, as indicated by similar T_FH_ cell, germinal center or class-switched B cell or plasma cell population counts in VP0 mRNA and control immunized mice (Fig. S2).

**Figure 3.**
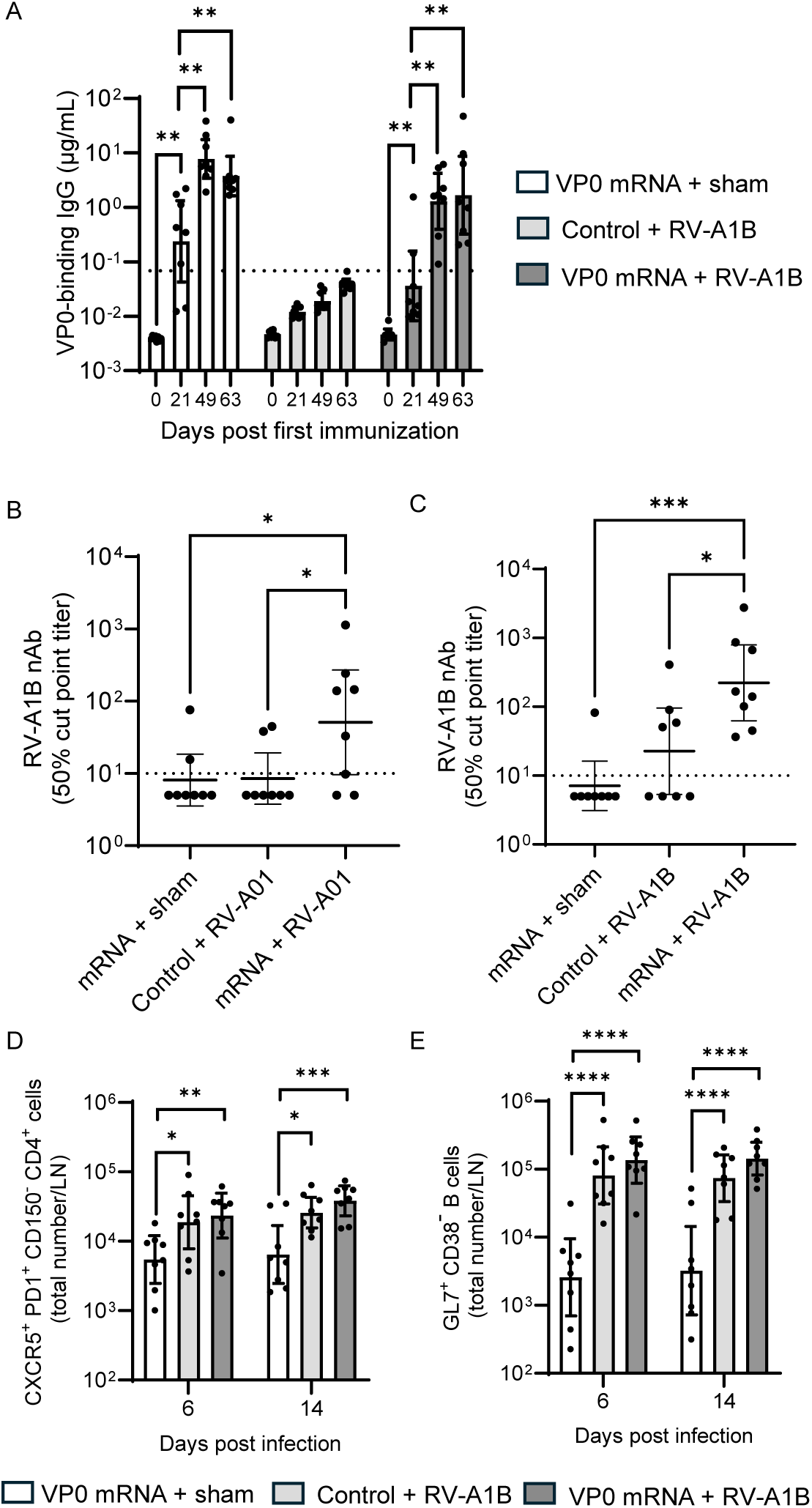
RV-A16 VP0 mRNA immunization evokes limited neutralizing antibody production to a heterotypic RV-A1B infection. Mice were immunized twice with RV-A16 VP0 mRNA or control 21 days apart. 28 days after the final immunization mice were infected intranasally with live RV-A1B or were sham-infected. Blood was taken on study days Day 0, 21, 49, 55 (6 days post infection) and 63 (14 days post-infection). The concentration of serum VP0-binding total IgG was determined at each timepoint in RV-A1B or sham infected animals (A). The titer of RV-A1B neutralizing serum antibodies was determined on day 6 (B) and 14 (C) post-infection. Mediastinal lymph node T_FH_ cell (D) and germinal centre B cell (E) abundance on Day 6 and 14 post-infection was determined by flow cytometry. **** p<0.0001, *** p<0.001, ** p<0.01, * p<0.05 (ANOVA). Graphs show geomeans with 95% CI. Each point represents an individual mouse.

RV-A16 VP0 mRNA immunization boosted the titer of RV-A1B nAbs at Day 6 post infection (Fig. 3B), which were higher again on Day 14 (Fig. 3C), but the titers were lower than those observed with rVP0 immunization (Fig. 2C). Consistent with a lower effect on post-infection RV-A1B nAb titers, there was no difference in the abundance of mediastinal (lung-draining) lymph node T_FH_ cells or germinal centre B cells between VP0 mRNA or control immunized mice at either of the post infection timepoints measured (Fig. 3D & E), or class-switched B cells or plasma cells (data not shown).

### Heterologous RV infection boosts broadly cross-reactive T cell responses in mice previously vaccinated with RV-A16 VP0 mRNA

Splenocytes isolated on Day 6 and Day 14 post infection were stimulated with peptide pools corresponding to full-length VP0 from a much larger and more diverse set of twenty different RV-A, RV-B and RV-C subtypes than the protein vaccination study (Fig. 4A), including RV-A16 (vaccine homologous) and RV-A1B (heterologous challenge), and IFN-γ secretion determined by ELISpot.

**Figure 4.**
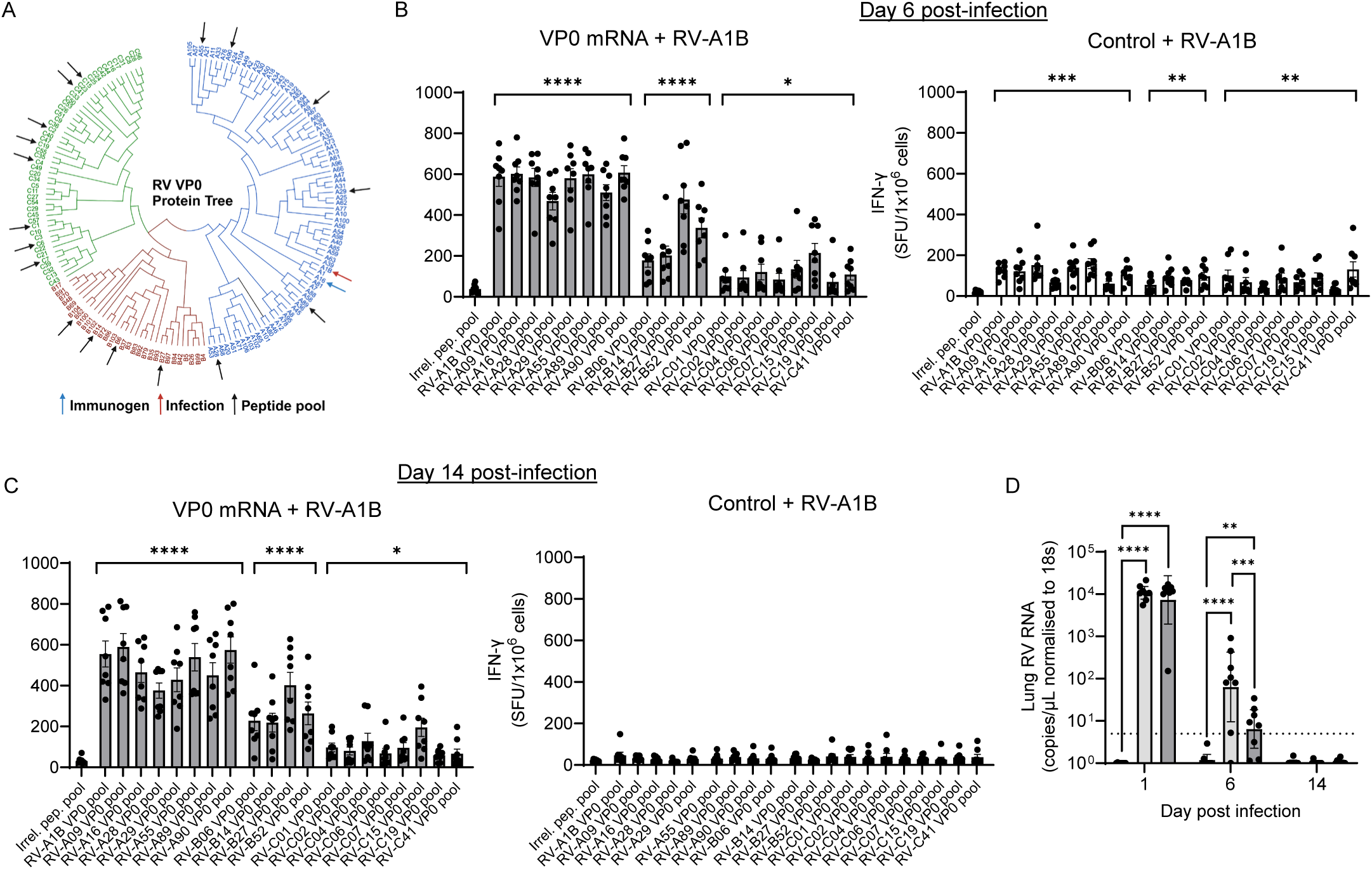
RV-A16 VP0 mRNA immunization creates cross-reactive cellular anti-RV immunity and enhances clearance of heterotypic virus. Mice were immunized twice with RV-A16 VP0 mRNA 21 days apart. 28 days after the final immunization mice were infected intranasally with live RV-A1B. Spleens were harvested on study days 55 (Day 6 post-infection) and Day 63 (Day 14 post-infection), splenocytes prepared and stimulated with peptide pools corresponding to full-length VP0 (15-mers with 11 amino acid overlap) from twenty different RV-A, B and C strains (A) or a negative control irrelevant peptide pool (human HLA class I Ig-like C1 type domain). IFN-γ-secreting cells from control or RV-A16 VP0 mRNA immunized mice on day 6 (B) and day 14 (C) post-infection were quantified by ELISpot. Lungs were harvested on days 1, 6, and 14 post-infection and viral RNA abundance determined by RT-qPCR (D). Dotted line indicates LLOQ. **** p<0.0001, *** p<0.001, ** p<0.01, * p<0.05 (ANOVA) compared to irrelevant peptide pool. (B – C) Graphs show mean with SEM; (D) Graph shows geomeans with 95% CI. Each point represents an individual mouse.

Strikingly, on Day 6 after RV-A1B infection, all RV-A and RV-B peptide pools tested stimulated a strong IFN-γ response in splenocytes from RV-A16 mRNA vaccinated mice (Fig. 4B), with strong responses persisting to Day 14 (Fig. 4C). However, the responses to RV-C VP0 peptide pools, although statistically significant, were considerably lower. RV-A1B infection also triggered a modest IFN-γ response in splenocytes from adjuvant only immunized mice against all of the RV-A VP0 peptide pools tested (Fig. 4B). However, this response in mock-immunized mice was transient (i.e. disappeared by Day 14 post infection) (Fig. 4C) and of lower magnitude than in RV-A16 VP0 mRNA vaccinated mice and hence likely reflective of primary T cell activation caused by infection.

In mice, RV-A1B infection is self-limiting and of short duration. On Day 1 post-infection viral RNA was detected in similar quantities in mice that had been previously immunized with RV-A16 VP0 mRNA vaccine or with adjuvant only control (mock-immunized controls) but not in sham-infected controls (Fig. 4D). By day 6 post-infection, viral infection was being cleared from lungs as indicated by viral genome counts having decreased in all infected mice. However, viral genome counts were significantly lower in RV-A16 VP0 mRNA immunized mice than in mock-immunized mice (approximately 30-fold), indicating accelerated virus clearance from the lung as result of RV-A16 VP0 mRNA vaccination. By day 14 after infection, mice in both RV-A16 VP0 mRNA- and control-immunization groups had cleared the infection.

### Sustained CD4^+^ and CD8^+^ T cell infiltrates, rich in effector cell subsets, are boosted in the airways of RV-A16 VP0 mRNA vaccinated mice following RV-A1B infection

In accordance with observations in humans, infection of mice with RV-A1B had a marked impact on recruitment of lymphocytes into the respiratory tract. Following virus challenge, lymphocyte infiltrates in the airways were significantly increased in all RV-A16 VP0 mRNA immunized mice, although in the early part of infection (Days 1 and 6) their absolute numbers and relative frequencies were similar to those in mock-immunized mice (Fig. 5A). By Day 14 post-infection, lymphocytes were significantly more abundant and made up a greater proportion of total airway leukocytes in RV-A16 VP0 mRNA immunized mice than in control mock immunized mice, suggesting a more sustained lymphocyte response in mice immunologically primed by VP0 during the viral clearance phase.

**Figure 5.**
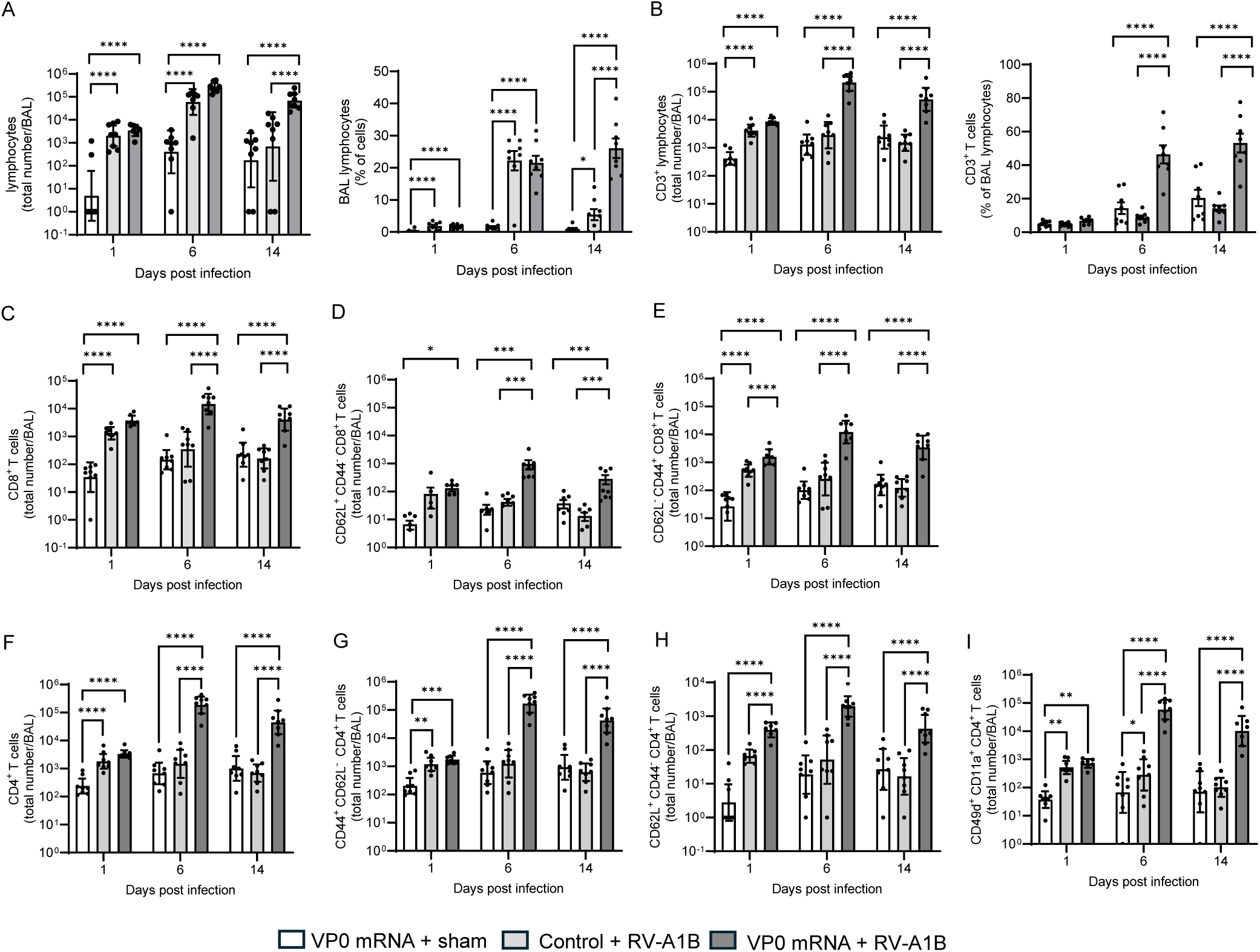
RV-A16 VP0 mRNA immunization promotes airway T cell expansion in response to heterotypic RV-A1B infection. Mice were immunized twice with RV-A16 VP0 mRNA 21 days apart. 28 days after the final immunization mice were infected intranasally with live RV-A1B or were sham-infected. BAL was harvested on study days 50 (Day 1 post-infection), 55 (Day 6 post-infection) and Day 63 (Day 14 post-infection), single cell suspensions prepared and T cell subset abundance and frequency, as indicated, determined by flow cytometry. **** p<0.0001, *** p<0.001, ** p<0.01, * p<0.05 (ANOVA). Graphs on log scale show geomeans with 95% CI, those on a linear scale show mean with SEM. Each point represents an individual mouse.

Total BAL CD3^+^ T cell counts were significantly higher in RV-A16 VP0 mRNA immunized mice at Day 6 and Day 14 post infection, compared to mock-immunized mice, which was also reflected in greater frequencies of CD3^+^ T cells as a proportion of total lymphocytes in the same mice (Fig. 5B). CD8^+^ T cell numbers were increased in BAL at Day 1 post-infection in all virus-challenged mice as compared to sham-challenged mice (Fig. 5C). However, by Day 6 post-infection CD8^+^ T cell numbers had further increased in VP0 mRNA immunized mice and remained elevated on Day 14 post-infection, while in mock-immunized control mice, CD8^+^ T cells numbers successively contracted from Day 1 to Day 6 and Day 14 post infection, reaching levels similar to those observed in sham infected mice at the same time points. The increase in CD8^+^ T cell counts in BAL from VP0 mRNA immunized mice post infection was associated with an increase in number of both naïve (CD62L^+^ CD44^-^) (Fig. 5D) and effector (CD62L^-^CD44^+^) (Fig. 5E) CD8^+^ T cell subsets, with effector CD8^+^ T cell numbers being approximately 1 order of magnitude higher than naïve CD8^+^ T cells.

The trends observed in the airway CD4^+^ T cell compartment were similar to those observed with the CD8^+^ T cell compartment, with BAL from VP0 mRNA immunized and from control immunized mice exhibiting higher CD4^+^ T cell counts than sham infected mice at Day 1 post infection, but RV-A16 VP0 mRNA immunized mice having much higher CD4^+^ T cell counts than either control group at Day 6 and 14 post infection (Fig. 5F). Of note, the number of airway CD4^+^ T cells from RV-A16 VP0 mRNA immunized mice on Day 6 and 14 post infection was ∼1 log higher than the number of CD8^+^ T cells.

The increase in CD4^+^ T cell counts in BAL from VP0 mRNA immunized mice at Day 6 and 14 post-infection was driven primarily by effector (CD44^+^ CD62L^-^) CD4^+^ T cells (Fig. 5G), although numbers of naïve (CD62L^+^ CD44^-^) CD4^+^ T cells (Fig. 5H) also significantly increased in these mice following viral challenge. It has been reported that CD49d^+^ CD11a^+^ CD4^+^ T cells are actively involved in response to viral infection and vaccination and represent antigen-specific CD4^+^ T cell subsets^21,22^. The abundance and frequency of airway CD49d^+^ CD11a^+^ CD4^+^ T cells at Day 6 and 14 post-infection was much greater in RV-A16 VP0 mRNA immunized than in mock-immunized control mice or in sham infected mice (Fig. 5I).

Put together, these findings suggest that clearance of RV-A1B from the respiratory tract in VP0 mRNA-immunized, but not in mock-immunized mice, was accompanied by robust infiltration of activated CD4^+^ and CD8^+^ T cells.

### RV-A16 VP0 mRNA vaccination induces sustained infiltration of CD8^+^ and CD4^+^ effector lymphocytes in the lungs of RV-A1B infected mice

Having dissected the cellular immune response to RV-A1B infection in the airways (BAL) of RV-A16 VP0 mRNA immunized mice, we next profiled the leukocyte infiltrate in lung tissue collected from the same mice.

The kinetics of T cell infiltration observed in the lung differed from that observed in the airways. In the airways there was evidence of a rapid CD4^+^ and CD8^+^ T cell infiltration at Day 1 post infection in both VP0 mRNA immunized and control immunized mice, a response which was subsequently amplified in VP0 mRNA immunized mice but not in mock-immunized mice at later timepoints. In the lungs, there was a slower infiltration of T cells following infection, with similar numbers of T cells on Day 1 post-infection in mice immunized with VP0 mRNA, mock-immunised mice or sham-infected mice (Fig. S3A). However, by Day 6 post-infection and to a lesser extent by Day 14 post-infection, the number of T cells was significantly greater in lungs harvested from RV-A16 VP0 mRNA immunized mice than from mock-immunized mice, or sham-infected mice. This finding was reflected in the frequency of T cells as a proportion of total lymphocytes observed in lung tissue (Fig. S3B), whereby Day 6 and Day 14 post-infection the percentage of T cells in the infiltrating lymphocyte population reached >20% in the lungs of most VP0 mRNA immunized mice but remained ∼10% in control mock-immunized or sham-infected mice.

Lung CD8^+^ T cell counts after infection also reflected those observed with total T cells, with VP0 mRNA vaccinated mice exhibiting greater CD8^+^ T cell counts in lung than any of the control groups on both Day 6 (Fig. 6A). The increase in CD8^+^ T cells post infection, was driven by expansion of both naïve (Fig. 6B) and effector (Fig. 6C) cells, with a relatively bigger increase in effector cells in VP0 mRNA immunized mice compared to mock-immunized or sham-infected controls.

**Figure 6.**
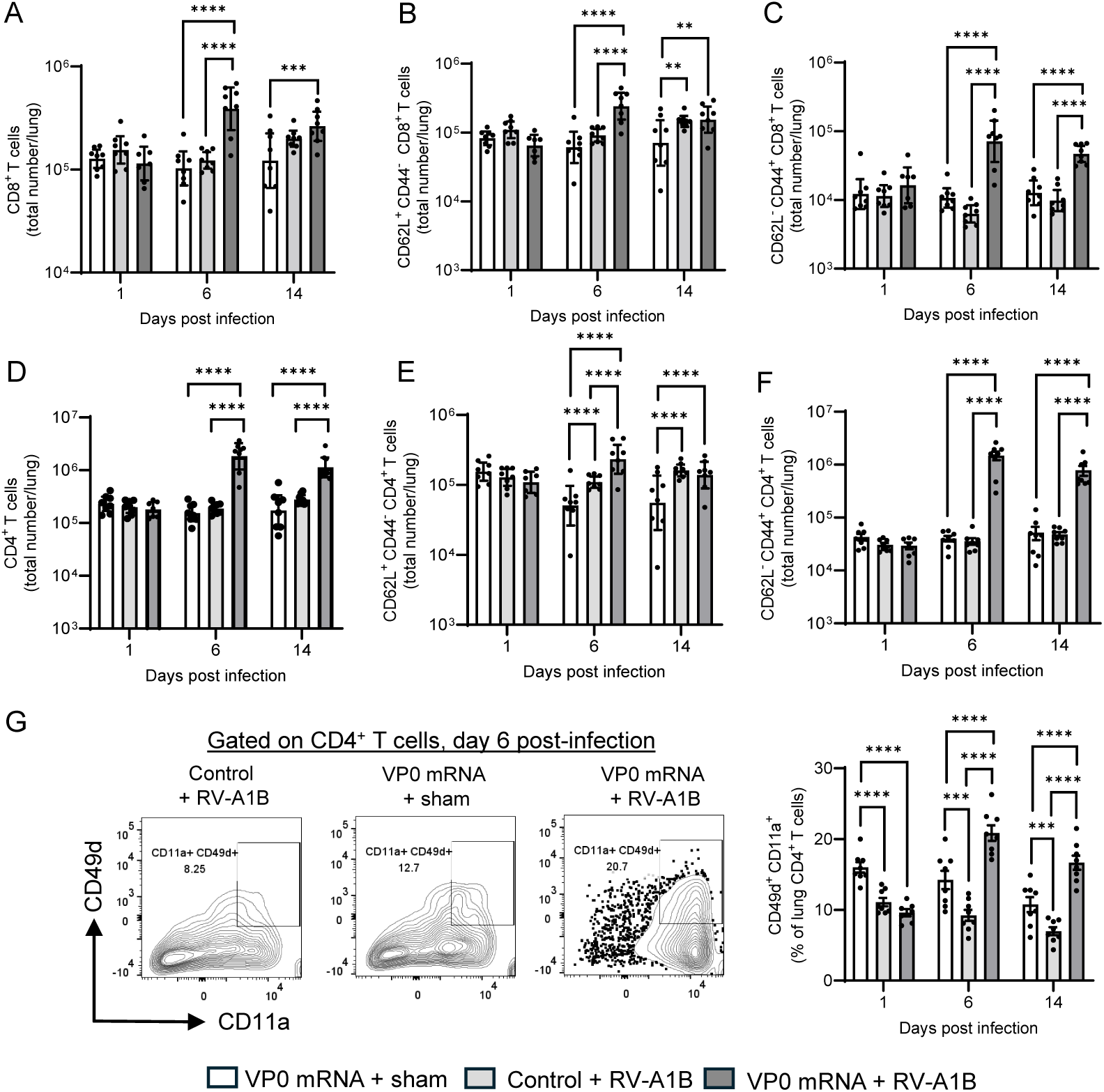
RV-A16 VP0 mRNA immunization promotes lung T cell activation and expansion in response to heterotypic RV-A1B infection. Mice were immunized twice with RV-A16 VP0 mRNA 21 days apart. 28 days after the final immunization mice were infected intranasally with live RV-A1B. Lungs were harvested on study days 50 (Day 1 post-infection), 55 (Day 6 post-infection) and Day 63 (Day 14 post-infection), single cell suspensions prepared and T cell subset abundance and frequency, as indicated, determined by flow cytometry. **** p<0.0001, *** p<0.001, ** p<0.01, * p<0.05 (ANOVA). Graphs on log scale show geomeans with 95% CI, those on a linear scale show mean with SEM. Each point represents an individual mouse.

Lung CD4^+^ T cells on Day 6 and 14 post infection showed the same trend for increased numbers in VP0 mRNA immunized mice, but not in mock-immunized controls or sham-infected mice (Fig. 6D). However, in VP0 mRNA vaccinated mice, CD4^+^ T cells made up a greater proportion (*c.a.* 80%) of the total CD3^+^ T cell infiltrate found in lungs post infection than CD8^+^ T cells (*c.a.* 20%) (Fig. S3C and S3D). The CD4^+^ cell infiltrate was so dominant in VP0 mRNA vaccinated mice post-infection, that it greatly skewed the frequency of CD8^+^ T cells as a proportion of total T cells, as compared to control mock-immunized or sham infected mice (Fig. S3D).

As in the airways, both naïve (Fig. 6E) and effector (Fig. 6F) CD4^+^ T cell numbers increased in the lung after RV infection in VP0 mRNA immunized mice, however, the expansion of effector CD4^+^ T cells was much greater than the expansion of naïve cells. This was reflected in the frequencies of naïve and effector CD4^+^ T cells, with >60% of CD4^+^ T cells being comprised of effector cells (Fig. S3E) and <20% of naïve cells (Fig. S3F). This is in direct contrast to observations in lungs from control immunized mice post infection, where naïve CD4^+^ T cells were the dominant phenotype (Fig. S3E). As in the airways, there was also a marked increase in lung infiltrates of antigen-specific CD49d^+^ CD11a^+^ CD4^+^ T cells following infection of VP0 mRNA immunized mice, compared to mock-immunized controls (Fig. 6G).

### Vaccination with RV-A16 VP0 mRNA generates a tissue resident memory (T_RM_) response following RV-A1B infection

Tissue resident memory T (T_RM_) cells in the airways, which constitute the entry point for respiratory infections, are vital contributors to protective immunity. We therefore measured the abundance and frequency of CD103^+^ CD69^+^ cells among CD4^+^ or CD8^+^ T cell populations in the BAL to determine if RV-A16 VP0 mRNA immunization primed animals for a rapid peripheral T_RM_ cell response to subsequent RV infection.

RV-A1B infection triggered marked expansion of both CD4^+^ and CD8^+^ T_RM_ lymphocyte populations in the airways of VP0 mRNA immunized but not mock-immunized mice, as reflected by significantly higher total counts of CD8^+^ (Fig. 7A) and CD4^+^ (Fig. 7B) T_RM_ cells in the BAL. Of note, following infection in VP0 mRNA immunized mice, CD8^+^ T_RM_ made up a greater proportion of all BAL CD8^+^ T cells (*c.a*. 40%) (Fig. 7C) than CD4^+^ T_RM_ did of all BAL CD4^+^ T cells (*c.a*.15%) (Fig 7D), as observed on Days 6 and 14 post-infection.

**Figure 7.**
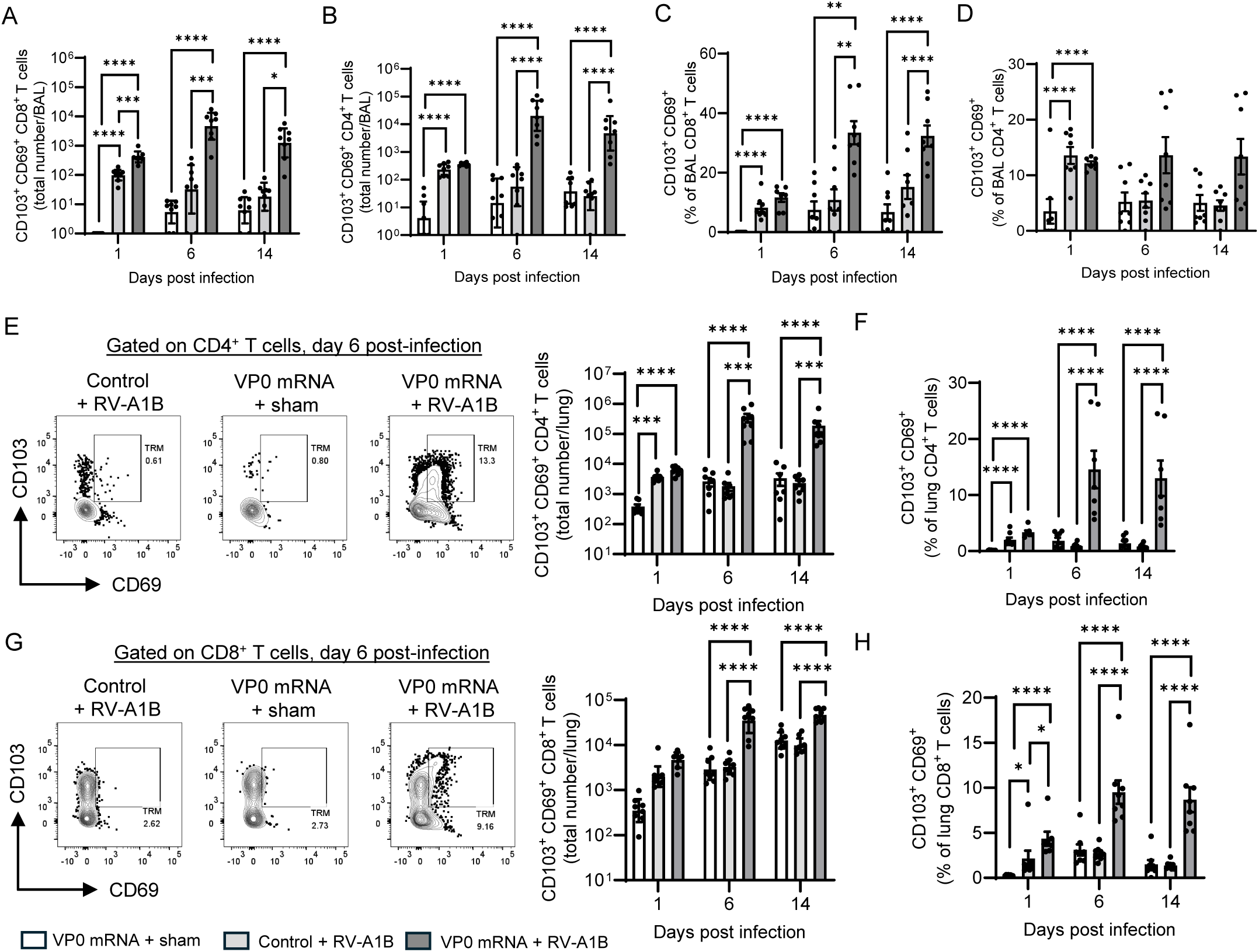
RV-A16 VP0 mRNA immunization boosts tissue resident T cell memory in the airways and lungs in response to heterotypic RV-A1B infection. Mice were immunized twice with RV-A16 VP0 mRNA 21 days apart. 28 days after the final immunization mice were infected intranasally with live RV-A1B or were sham-infected. BAL and lung lobes were harvested on study days 50 (Day 1 post-infection), 55 (Day 6 post-infection) and Day 63 (Day 14 post-infection), single cell suspensions prepared and T_RM_ abundance and frequency, as indicated, determined by flow cytometry. **** p<0.0001, *** p<0.001, ** p<0.01, * p<0.05 (ANOVA). Bar charts on log scale show geomeans with 95% CI, those on a linear scale show mean with SEM. Each point represents an individual mouse.

In the lungs also, there were significant increases in the frequency and abundance of CD4^+^ (Fig. 7E-F) and CD8^+^ (Fig. 7 G-H) T_RM_ cells in RV-A16 VP0 mRNA immunized mice but not in mock-immunized controls after RV-A1B infection, indicating that VP0 mRNA immunization primes for peripheral immune memory to heterotypic virus infection.

### RV-A16 VP0 mRNA vaccination produces a strong local VP0-specific cross-reactive Th1 response in the lungs of RV-A1B infected mice

Having demonstrated that robust effector T cell expansion is triggered by RV-A1B infection in the lungs and airways of RV-A16 VP0 mRNA immunized mice, we went on to explore the antigen-specificity and phenotype of these cells. To this effect, lung cells were isolated from mice that had been infected with RV-A1B/sham after immunization with RV-A16 VP0 mRNA vaccine or mock vaccine.

Lung cells harvested on Day 1, 6 and 14 post-infection, were stimulated *ex vivo* with peptide pools corresponding to full length VP0 protein from the infecting RV-A1B strain or the heterologous RV-A09 strain and stained for Th1 (CD4, T-bet, IFN-γ) or Th2 (CD4, Gata3 & IL-4) markers to determine the frequency and phenotype of reactive cells. There was increased abundance of Th1 cells following stimulation with either RV-A1B or RV-A09 VP0 peptides in lung cells from VP0 mRNA immunized/RV-A1B infected mice compared to controls (Fig. 8A and 8C). In contrast, there was no concomitant increase in Th2 cells (Fig. 8B and 8D), demonstrating that systemic VP0 mRNA immunization primes a local Th1-biased response to infection in the lungs. Furthermore, that RV-A09, an RV-A subtype distantly related RV-A16 and RV-A1B, stimulated lung cells from VP0 mRNA immunized/RV-A1B infected mice further demonstrates that VP0 mRNA immunization evokes a cross-reactive cellular immune response to subsequent challenge. Direct comparison between the proportion of Th1 and Th2 cells following RV-A1B or RV-A09 peptide stimulation confirmed a predominantly Th2 response in unvaccinated animals (Fig. 8E and 8G) and a Th1-dominated response in lung cells from VP0 mRNA immunized/RV-A1B infected mice (Fig. 8F and 8H).

**Figure 8.**
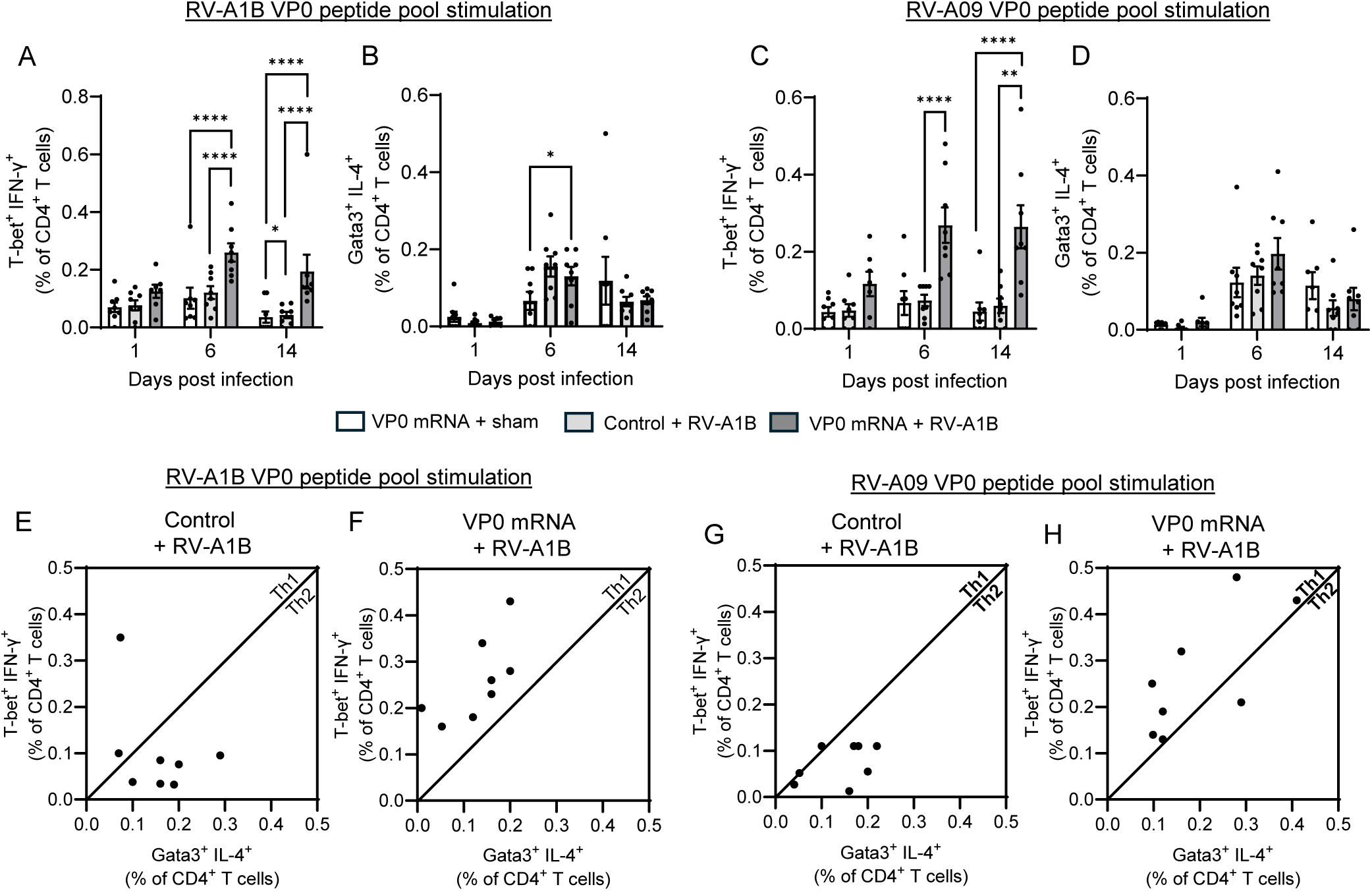
RV-A16 VP0 mRNA immunization evokes a Th1 polarized cross-reactive immunity. Mice were immunized twice with RV-A16 VP0 mRNA 21 days apart. 28 days after the final immunization mice were infected intranasally with live RV-A1B. Lungs were harvested on days 1, 6 and 14 post-infection, single cell suspensions prepared and stimulated with peptide pools corresponding to full length VP0 from RV-A1B (A, B, E & F) or RV-A09 (C, D, G & H). The frequency of T-bet^+^ IFN-γ^+^ (Th1) and Gata3^+^ IL-4^+^ (Th2) CD4^+^ T cells, as indicated, was determined by flow cytometry (A-D). (E-H) frequency of Th1 vs Th2 cells from each mouse. **** p<0.0001,, ** p<0.01, * p<0.05 (ANOVA). Bar graphs show means with SEM. Each point represents an individual mouse.

## Discussion

RV infection is the primary cause of exacerbations of chronic lung diseases, leading to substantial morbidity and mortality. To date, RV vaccine development has been hampered by the large number of circulating RV subtypes and the high degree of antigenic diversity among subtypes. We have explored the potential of RV-A16 VP0 as a RV vaccine immunogen in adjuvanted protein subunit and mRNA formats and demonstrated that RV-A16 VP0 induces broadly cross-reactive cellular immunity that protects mice from subsequent heterotypic RV challenge. Furthermore, we have identified differences between the immune responses to RV-A16 VP0 when administered as adjuvanted protein or mRNA.

There are currently no approved vaccines or therapeutics that directly target RV to protect clinically vulnerable COPD patients from the most common cause of acute exacerbations. Small molecule inhibitors of RV replication are in clinical testing (NCT06149494, NCT05398198), but these would be administered after infection, which may limit their effectiveness. Therefore, the holy grail remains a safe and effective vaccine.

The induction of rapid antigen-specific cell mediated immunity by viral vaccines has been exemplified by the success of the SARS-CoV-2 vaccines, specifically by those that employ mRNA technology^23^. Furthermore, protective immunity mediated by T cells has been demonstrated against influenza H1N1 in neutralizing antibody seronegative subjects^24^. We hypothesized that both adjuvanted recombinant RV-A16 VP0 protein subunit and mRNA vaccine platforms would evoke broadly reactive cellular immunity to diverse RV subtypes, which could underpin the mechanistic basis of broad protection in humans.

In mice subsequently infected with heterotypic RV-A1B subtype, both adjuvanted protein and mRNA vaccine platforms evoked Th1 polarized cellular immunity that cross-reacted strongly with VP0 peptide pools from diverse RV-A and RV-B subtypes, and significantly, but to a lesser extent against RV-C subtypes. The response to infection in immunized mice was associated with expansion of CD4^+^ and CD8^+^ T cell populations in the airways and lungs, in particular effector and antigen-specific T cells. The enhanced T cell response observed in immunized mice was associated with accelerated virus RNA clearance from the lungs.

A critical component of T cell mediated immunity to viral pathogens are long-lived T_RM_ cells at the peripheral sites of infection^25,26^, in the case of RV infections - the airways and lungs. Here we have demonstrated that mice immunized with RV-A16 VP0 mRNA have significantly more T_RM_ cells in both the airways and lungs following infection, indicating that the vaccine primes both central and peripheral immunity. In patients, we hypothesise that priming of cross-reactive T_RM_ immunity in the airways and lungs will overcome the innate immune dysfunction often associated with chronic lung diseases such as COPD^27^ and effectively clear the RV infection before it causes exacerbation.

An RV vaccine based upon a capsid protein from a single subtype is unlikely to evoke *de novo* cross-neutralizing humoral immunity, as has been evidenced by multiple preclinical and clinical studies^13–15,28^ . However, vaccine priming of cross-reactive CD4^+^ T cell anti-RV immunity should accelerate the humoral response to a subsequent heterotypic infection by providing help to B cells. Indeed, in mice immunized with RV-A16 rVP0 we detected RV-A1B nAbs by day 6 post-infection, whereas in control immunized mice nAbs were only detected on Day 14. By this mechanism, cross-reactive cellular immunity can accelerate and boost neutralizing antibody responses to subsequent infection.

In comparing the two vaccine platforms, the biggest difference we observed was in the humoral response. Antigen-specific IgG titer was lower in RV-A16 VP0 mRNA immunized animals than rVP0 immunized and was associated with reduced lymph node cellular responses. Furthermore, the neutralizing antibody response to subsequent infection was less in VP0 mRNA immunized mice. Overall, this suggests that the mRNA format used in this study did not evoke potent B cell activation. The differences between formats could be accounted for by the potent B cell stimulating adjuvant combination used with the rVP0 vaccine and/or the lack of a specific adjuvant/lipid nanoparticle, which has adjuvant properties^29^, in the mRNA formulation. Both the formulation and timing of delivery for mRNA vaccines has been shown to impact the immune response; further work to optimise RV-A16 VP0 mRNA delivery may provide a more comparable response to rVP0 immunisation.

While RV-A16 VP0 immunization elicits cellular immunity that cross-reacts with subtypes from all 3 species of RV, the degree of cross-protection against RV-C subtypes was comparatively lower than that against RV-A and RV-B subtypes. Therefore, RV-A16 may not be the optimal VP0 sequence to generate truly cross-protective immunity to RV subtypes most often linked to inflammatory disease exacerbation in humans and a different and/or multiple VP0 sequences may provide greater breadth of protection.

## Methods

### Animals

All protocols for mouse experimental work were approved by the UK Home Office and performed under license PP0352637 granted 17-Feb-21. Female specific pathogen free (SPF) C57BL/6J mice were purchased from Charles River Laboratories (Harlow, UK) at 6-8 weeks old and housed in SPF conditions at the Royal Veterinary College (Camden, UK). Procedures and animal care conformed to the Animals (Scientific Procedures) Act 1986 guidelines and were conducted under local Animal Welfare and Ethical Review Body (AWERB) guidance at the Royal Veterinary College, Camden (UK).

### Immunogens

Full-length RV-A16 VP0 protein was produced in *E.Coli* using methods previously described^20^. Fully pseudouridine substituted full-length RV-A16 VP0 mRNA was produced by TriLink BioTechnologies (US). RV-A16 VP0 protein was adjuvanted with 6.3 μg CpG ODN 1826 (InvivoGen, US) and Incomplete Freund’s Adjuvant (Merck, UK). RV-A16 VP0 mRNA was formulated for injection using *in vivo*-jetRNA® and mRNA buffer (both PolyPlus, FR).

### Animal models of immunization and RV infection

Mice were immunized s.c. (lower abdominal quadrant) with 10 μg RV-A16 VP0 protein or i.m. (thigh muscle) with 5 μg RV-A16 VP0 mRNA. RV-A16 VP0 protein was adjuvanted with 6.3 μg CpG ODN 1826 (InvivoGen, US) and Incomplete Freund’s Adjuvant (Merck, UK); RV-A16 VP0 mRNA was formulated in *in vivo*-jetRNA® reagent nanoparticles and mRNA buffer (both PolyPlus, FR).

For studies with subunit vaccines, mice were immunized s.c. with RV-A16 rVP0 twice, 21 days apart; for studies with mRNA vaccines, mice were immunized i.m. with RV-A16 VP0 mRNA twice, 21 days apart. Inguinal lymph nodes were harvested 10 days after the second immunization, spleens and mediastinal lymph nodes were harvested 6 and 14 days post-infection, blood samples were taken pre-immunization on Day 0 and Day 21 and pre-infection on Day 49, in addition to terminal bleeds.

RV-A1B was grown, purified and titrated for *in vivo* use as described previously^30^. Mice were dosed intranasally with 10^7^ TCID_50_ RV-A1B in 50 μL, administered under light anesthesia.

### Sample collection and processing

Prior to termination, serial blood samples of ≤ 50 μL were collected from the tail vein using Hirschmann capillary tubes (VWR, UK) and centrifuged to isolate serum. At study endpoints, mice were euthanized with an overdose of anesthetic delivered i.p. and exsanguinated via the jugular vein. Whole blood was collected using Hirschmann capillary tubes and centrifuged to isolate serum.

To harvest BAL, the lungs were flushed with 3 x 0.5 mL sterile PBS (Gibco, UK) via a cannula placed into the trachea. Whole BAL samples were separated into cells and supernatants by centrifugation.

Inguinal and mediastinal lymph nodes and spleens were harvested into R10 media (RPMI containing 10% FBS and 1% P/S (all Gibco)). Single cells were isolated from each tissue by passing through 100 µm EASYstrainer (Greiner Bio-One, UK). Red blood cells in the spleen isolates were lysed by incubation with ACK buffer (Gibco).

Superior and post-caval lung lobes were harvested and snap frozen. All remaining lobes were finely diced and incubated at 37°C for 30 minutes in R10 media containing 0.14 WU / mL Liberase 3 and 0.05 mg / mL DNAse I (both Sigma-Aldrich, UK), then processed into single cell suspensions as outlined above. Red blood cells were lysed by incubation with ACK buffer.

### Semi-quantitative RV-A16 VP0-specific IgG ELISA

Nunc™ MaxiSorp™ 96-well ELISA microplates (Fisher Scientific) were coated with 25 ng RV-A16 VP0 antigen (WuXi Biologics, China) or 50 ng mouse IgG antibody (R&D Systems, US) for the standard curve. Plates were subsequently blocked using PBS 2% BSA (A7906, Sigma Aldrich) then incubated with serum samples diluted in PBS 2% BSA (Sigma Aldrich), or standards for 2 hours at room temperature. The standard curve was created using mouse IgG ELISA standard (Fisher Scientific) at a top concentration of 100 μg/mL followed by a 1 in 3 dilution series. Bound antibody was detected using goat anti-mouse biotinylated IgG (R&D Systems, US) at 5 μg/mL, followed by streptavidin-HRP at 1 μg/mL. Reactions were developed using 1-Step™ TMB substrate (Thermo Fisher, UK) per well, followed by 50 uL 0.18M H_2_S0_4_ (Sigma Aldrich) and read at OD 450 nm using a SpectraMax iD3 analyzer (Molecular Devices, US).

### Flow Cytometry

Lung, lymph node and BAL single cell preparations were counted using trypan blue exclusion staining (Gibco): 1-2 million cells were used per sample for surface stains, 3-4 million for intracellular/intranuclear stains. To analyze cytokine and transcription factor expression, lung cells were stimulated prior to staining with 16 μg/ mL RV-A1B or RV-A09 VP0 peptides (Mimotopes, AU) in the presence of 10 μg/ mL brefeldin A (Enzo, US) for 4 hours at 37°C.

All cells were washed with PBS and then incubated with fixable violet LIVE/DEAD™ stain (Molecular Probes, UK) following manufacturer’s instructions. The following antibodies were used to stain surface markers, diluted in PBS 2 % FBS (both Gibco) 2 mM EDTA (Invitrogen, UK) containing TruStain FcX™ anti-CD16/32 (BioLegend) at dilutions previously established by titration: anti-CD4 (RM4-5), anti-CD185 (L138D7), anti-PD-1 (RMP1-30), anti-CD150 (W19132B), anti-IgD (11-26c.2a), anti-CD19 (6D5), anti-GL7 (GL7), anti-CD38 (90), anti-CD90.2 (30-H12), anti-CD8 (53-6.7), anti-CD45R (RA306B2), anti-CD3ε (14502C11), anti-CD62L (MEL-14), anti-CD44 (IM7), anti-CD11a (M17/4), anti-CD69 (H1.2F3) (all BioLegend, UK), anti-CD45R (RA306B2), anti-CD49d (R1-2), anti-CD103 (2E7) (all eBioscience, UK). Surface stains were fixed with PBS 1% PFA (Sigma Aldrich). For intracellular and intranuclear stains, cells were fixed with Foxp3/ Transcription Factor Staining buffer kit (eBioscience), then incubated with the following antibodies diluted in permeabilization buffer (eBioscience): anti-T-bet (4B10), anti-IL-4 (11B11), anti-IL-10 (JES5-16E3) (all BioLegend), anti-Foxp3 (FJK-16s), anti-Gata-3 (TWAJ), anti-IFNγ (XMG1.2) (all eBioscience). Samples were acquired using a 3-laser BD Fortessa and analyzing using FlowJo™ Software (BD Life Sciences).

### RV-A1B neutralising antibody titrations

Mouse serum was diluted in 5×10^4^ TCID_50_ per mL RV-A1B following a 1/10^0.5^ dilution series to create an 8-point curve then incubated for 1 hour prior to plating with H1HeLa cells (ATCC, UK) at 3×10^5^ cell per mL in a 96-well TC-treated microplate (VWR). Assays were incubated at 37°C for 72 hours. Following incubation, monolayers were stained with 0.1% crystal violet solution (Sigma-Aldrich) then solubilized in a PBS 1% SDS solution (Fisher BioReagents, UK). Plates were read at OD 560 nm using a SpectraMax iD3 analyser (Molecular Devices, US).

### IFN-γ and IL-5 ELISpots

MultiScreen® 96-well plates (Sigma Aldrich) were coated with 0.5 μg/mL anti-IL-5 antibody or 5 ug/mL IFN-γ (both BD Biosciences) overnight at 4°C, then blocked with R10 medium. 10^6^ splenocytes were plated per well and incubated overnight with one of the following stimuli: medium alone, 50 ng/mL PMA (Invivogen) 500 ng/mL ionomycin (Stemcell Technologies, UK), 25 μg/mL RV-A16 VP0 (WuXi Biologics), 16 μg/mL RV VP0 peptides (Mimotopes) or 16 μg/mL irrelevant peptide (JPT Peptides, DE). Following incubation, cells were lysed with diH_2_O. Bound cytokine was detected using 1 μg/mL biotinylated anti-IL-5 or 2 µg/mL biotinylated anti-IFN-γ (both BD Biosciences) followed by 15 μg/mL ExtrAvidin® Alkaline Phosphatase (Sigma-Aldrich). Spots were developed by incubation with SIGMAFAST™ BCIP®/NBT tablets (Sigma-Aldrich) made as per manufacturers’ instructions. Plates were allowed to air dry and then analyzed using an AID Classic ELISpot Reader (AID, DE).

### Rhinovirus RT q-PCR

Total RNA was extracted from the superior lung lobe using RNeasy mini kits; 2 μg RNA was converted to cDNA using Omniscript® Reverse Transcription kit (both Qiagen, NL). RT-qPCR reactions were set up in 96-well optical PCR plates (Fisher Scientific) using RV and 18s primers and probes as previously described^20^, combined with QuantiTect PCR mastermix (Qiagen) and extracted cDNA, along with an 8-point RV-A1B standard curve (10^7^ – 10^0^ copies per μL). RT-qPCR was performed using a QuantiStudio 5 Real Time PCR System and analyzed in QuantStudio Design and Analysis Software (both Applied Biosystems, UK).

### BAL cell differential counts

BAL cells were transferred to glass slides using a Cytospin™ 4 and stained once dry using Kwik-Diff™ Stain (both Epredia, US) following manufacturers’ instructions and viewed under an inverted light microscope (Leica Motic AE200) using a 40X objective. Two counts of one hundred cells were performed manually, identifying neutrophil, eosinophil, macrophage/ monocyte, and lymphocyte populations.

### mRNA VP0 western blot

HEK293 cells (Sigma Aldrich) were grown to 80% confluence in 24-well Cellstar® TC plates, then transfected with RV-A16 VP0 mRNA (TriLink BioTechnologies) using mRNA-Fect (RJH Biosciences, CA) following manufacturer’s instructions. Cells were incubated at 37°C 24 – 96 hours, then harvested into 2x SDS sample buffer (Invitrogen). Cell lysates were mixed with 10% beta-mercaptoethanol (Sigma-Aldrich), then denatured at 95°C for 5 minutes. 30 μL of HEK lysate, or 3 ng purified RV-A16 VP0, were combined with NuPage MES SDS running buffer and loaded into a 12% NuPage Bis-Tris polyacrylamide gel (both Invitrogen). 10 μL of See-Blue Plus 2 pre-stained marker (Invitrogen) was also loaded and gels were run at 50 mA for 30 mins then 180 mA for 90 minutes using a Mini Gel Tank (Invitrogen). Proteins were transferred onto 0.45 µm nitrocellulose membranes (Thermo Scientific) in NuPage transfer buffer (Invitrogen) containing 10 % methanol (Sigma-Aldrich) at 180 mA. Membranes were blocked using PBS 5% skim milk protein (Sigma-Aldrich), then probed with 2 µg/ mL anti-RV-A16 monoclonal antibody (Antibodies Online, US) followed by 0.2 µg / mL IgG (H+L) antibody HRP conjugate (Sigma-Aldrich). 2 mL ECL Plus substrate (Thermo Scientific) was added for 2min and membranes imaged using a Peqlab imager (Peqlab, UK).

## Data availability statement

All data generated or analysed during this study are included in this published article and its supplementary information files.

## References

1. Ritchie, A. I. et al. Pathogenesis of Viral Infection in Exacerbations of Airway Disease. Ann. Am. Thorac. Soc. 12, S115–S132 (2015).

2. Tan, K. S. et al. Respiratory viral infections in exacerbation of chronic airway inflammatory diseases: novel mechanisms and insights from the upper airway epithelium. Frontiers in Cell and Developmental Biology 8, 99 (2020).

3. Hewitt, R. et al. The role of viral infections in exacerbations of chronic obstructive pulmonary disease and asthma. Ther. Adv. Respir. Dis. 10, 158–174 (2016).

4. Friedlander, S. L. & Busse, W. W. The role of rhinovirus in asthma exacerbations. J. Allergy Clin. Immunol. 116, 267–273 (2005).

5. Venarske, D. L. et al. The Relationship of rhinovirus-associated asthma hospitalizations with inhaled corticosteroids and smoking. 193 (2006).

6. Hershenson, M. B. Rhinovirus-induced exacerbations of asthma and COPD. Scientifica 2013, 405876 (2013).

7. Seemungal, T. A., Hurst, J. R. & Wedzicha, J. A. Exacerbation rate, health status and mortality in COPD – a review of potential interventions. Int. J. Chronic Obstr. Pulm. Dis. 4, 203–223 (2009).

8. Cafferkey, J., Coultas, J. A. & Mallia, P. Human rhinovirus infection and COPD: role in exacerbations and potential for therapeutic targets. Expert Rev. Respir. Med. 14, 777–789 (2020).

9. George, S. N. et al. Human rhinovirus infection during naturally occurring COPD exacerbations. Eur. Respir. J. 44, 87–96 (2014).

10. Johnston, S. L. et al. Community study of role of viral infections in exacerbations of asthma in 9-11 year old children. BMJ 310, 1225–1229 (1995).

11. Palmenberg, A. C. Rhinovirus C, asthma, and cell surface expression of virus receptor CDHR3. J. Virol. 91, 10.1128/jvi.00072-17 (2017).

12. Gern, J. E. The ABCs of rhinoviruses, wheezing, and asthma. J. Virol. 84, 7418–7426 (2010).

13. Lee, S. et al. A polyvalent inactivated rhinovirus vaccine is broadly immunogenic in rhesus macaques. Nat. Commun. 7, 12838 (2016).

14. Glanville, N. & Johnston, S. L. Challenges in developing a cross-serotype rhinovirus vaccine. Curr. Opin. Virol. 11, 83–88 (2015).

15. McLean, G. R. Developing a vaccine for human rhinoviruses. Journal of vaccines & immunization 2, 16–20 (2014).

16. Appleyard, G. et al. Neutralization epitopes of human rhinovirus type 2. J. Gen. Virol. 71, 1275–1282 (1990).

17. Kan-o, K. et al. Differences in the spectrum of respiratory viruses and detection of human rhinovirus C in exacerbations of adult asthma and chronic obstructive pulmonary disease. Respir. Investig. 60, 129–136 (2022).

18. Bochkov, Y. A. et al. Rhinoviruses A and C elicit long-lasting antibody responses with limited cross-neutralization. J. Med. Virol. 95, e29058 (2023).

19. Choi, T. et al. Enhanced neutralizing antibody responses to rhinovirus C and age-dependent patterns of infection. Am. J. Respir. Crit. Care Med. 203, 822–830 (2021).

20. Glanville, N. et al. Cross-Serotype Immunity Induced by Immunization with a Conserved Rhinovirus Capsid Protein. PLoS Pathog. 9, e1003669 (2013).

21. McDermott, D. S. & Varga, S. M. Quantifying antigen-specific CD4 T cells during a viral infection: CD4 T cell responses are larger than we think. J. Immunol. 187, 5568–5576 (2011).

22. Christiaansen, A. F. et al. CD11a and CD49d enhance the detection of antigen-specific T cells following human vaccination. Vaccine 35, 4255–4261 (2017).

23. Painter, M. M. et al. Rapid induction of antigen-specific CD4+ T cells is associated with coordinated humoral and cellular immunity to SARS-CoV-2 mRNA vaccination. Immunity 54, 2133–2142.e3 (2021).

24. Sridhar, S. et al. Cellular immune correlates of protection against symptomatic pandemic influenza. Nature Medicine 19, 1305–1312 (2013).

25. Hassert, M. & Harty, J. T. Tissue resident memory T cells-A new benchmark for the induction of vaccine-induced mucosal immunity. Frontiers in Immunology 13, 5964 (2022).

26. Mueller, S. N. & Mackay, L. K. Tissue-resident memory T cells: local specialists in immune defence. Nat. Rev. Immunol. 16, 79–89 (2016).

27. Ritchie, A. I. et al. Pathogenesis of viral infection in exacerbations of airway disease. Annals of the American Thoracic Society 12, S115–S132 (2015).

28. Hamory, B. H., Hamparian, V. V., Conant, R. M. & Gwaltney, J. M. Human responses to two decavalent rhinovirus vaccines. J. Infect. Dis. 132, 623–629 (1975).

29. Alameh, M.-G. et al. Lipid nanoparticles enhance the efficacy of mRNA and protein subunit vaccines by inducing robust T follicular helper cell and humoral responses. Immunity 54, 2877–2892.e7 (2021).

30. Bartlett, N. W. et al. Mouse models of rhinovirus-induced disease and exacerbation of allergic airway inflammation. Nature medicine 14, 199–204 (2008).

