## Supplemental Figures for "A rhinovirus vaccine evokes T cell mediated cross-reactive immunity in a preclinical model of rhinovirus infection"

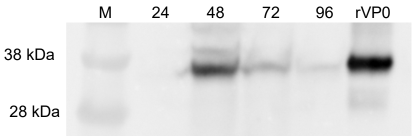


**Figure S1. RV-A16 VP0 mRNA is translated to full-length VP0 protein.** HEK293 cells were transfected with RV-A16 VP0 mRNA. Cell lysates were prepared 24, 48, 72 and 96 hours after transfection and RV-A16 VP0 expression assessed by Western blotting using an RV-A16 VP0 specific monoclonal antibody. M = marker, rVP0 = recombinantly expressed VP0 protein.

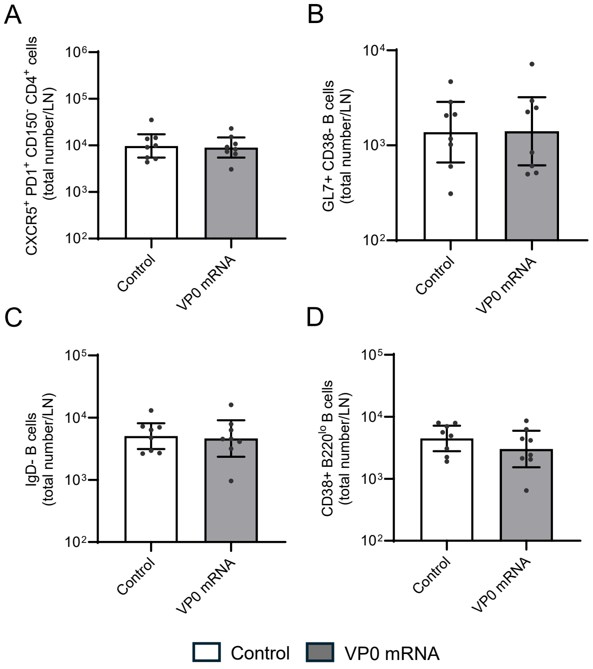


**Figure S2. RV-A16 VP0 mRNA immunization does not induce lymphoid responses**. Mice were immunized twice with RV-A16 VP0 mRNA or control 21 days apart. 10 days after the final immunization lymph nodes were harvested. The abundance of lymph node $T_{FH}$ (A), class-switched B cells (B), germinal center B cells (C) and plasma cells (D) were determined by flow cytometry. Each point represents an individual mouse.

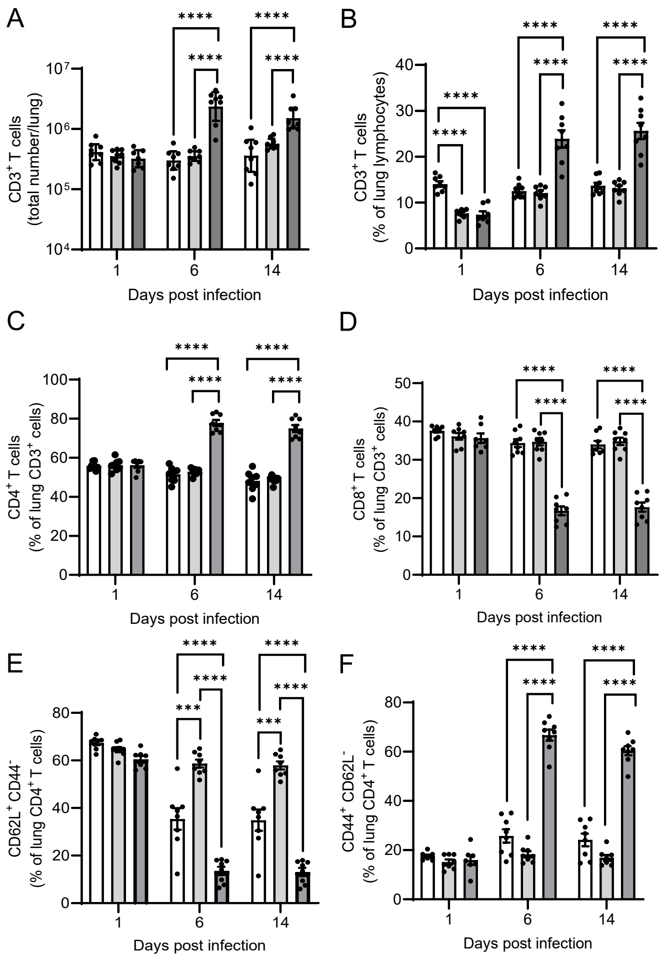


**Figure S3. RV-A16 VP0 mRNA immunization promotes lung T cell population expansion in response to heterotypic RV-A1B infection.** Mice were immunized twice with RV-A16 VP0 mRNA 21 days apart. 28 days after the final immunization mice were infected intranasally with live RV-A1B. Lungs were harvested on study days 50 (Day 1 post-infection), 55 (Day 6 post-infection) and Day 63 (Day 14 post-infection), single cell suspensions prepared and T cell subset abundance and frequency, as indicated, determined by flow cytometry. **** $p<0.0001$, *** $p<0.001$, ** $p<0.01$, * $p<0.05$ (ANOVA). Graphs on log scale show geomeans with 95% CI, those on a linear scale show mean with SEM. Each point represents an individual mouse.
